# Resting-State Electroencephalography and Magnetoencephalography in Migraine – A Systematic Review and Meta-Analysis

**DOI:** 10.1101/2024.08.02.606283

**Authors:** Paul Theo Zebhauser, Henrik Heitmann, Elisabeth S. May, Markus Ploner

**Author notes:** Corresponding author. Address: Department of Neurology, Technical University of Munich (TUM), Ismaninger Str. 22, 81675 Munich, Germany. (M. Ploner).

## Abstract

Magnetoencephalography/electroencephalography (M/EEG) can provide insights into migraine pathophysiology and help develop clinically valuable biomarkers. To integrate and summarize the existing evidence on changes in brain function in migraine, we performed a systematic review and meta-analysis (PROSPERO CRD42021272622) of resting-state M/EEG findings in migraine. We included 27 studies after searching MEDLINE, Web of Science Core Collection, and EMBASE. Risk of bias was assessed using a modified Newcastle–Ottawa Scale. Semi-quantitative analysis was conducted by vote counting, and meta-analyses of M/EEG differences between people with migraine and healthy participants were performed using random-effects models. In people with migraine during the interictal phase, meta-analysis revealed higher power of brain activity at theta frequencies (3-8 Hz) than in healthy participants. Furthermore, we found evidence for lower alpha and beta connectivity in people with migraine in the interictal phase. No associations between M/EEG features and disease severity were observed. Moreover, some evidence for higher delta and beta power in the premonitory compared to the interictal phase was found. Strongest risk of bias of included studies arose from a lack of controlling for comorbidities and non-automatized or non-blinded M/EEG assessments. These findings can guide future M/EEG studies on migraine pathophysiology and brain-based biomarkers, which should consider comorbidities and aim for standardized, collaborative approaches.

## 1 Background

Migraine affects over 1 billion people worldwide [1], substantially compromises the quality of life of those affected, and burdens healthcare systems and societies [2]. Despite substantial progress, the pathophysiology of this cyclic disease remains incompletely understood [2, 3]. Neuroimaging studies have uncovered widespread changes in brain structure and function in people with migraine [4]. These alterations have been observed in the brainstem, the hypothalamus, and an extended network of cortical areas [4]. Electroencephalography (EEG) and magnetoencephalography (MEG) are non-invasive, direct measures of neuronal activity that can further specify brain function in migraine. M/EEG studies during the resting state have shown both increased and decreased brain activity at different frequency bands [5–7]. Moreover, changes in functional connectivity and intrinsic brain network function have been observed [5, 8]. However, findings were not always replicable and partly conflicting. This inconsistency may be due to small sample sizes, different recording conditions, unstandardized data preprocessing, and diverse outcome measures. Moreover, findings might differ across the migraine cycle. Most studies have focused on M/EEG features in the interictal phase compared to healthy participants and associations of M/EEG features and disease severity. Other studies have recorded M/EEG during migraine attacks or the premonitory phase, and few studies have investigated variations across the migraine cycle [9]. Thus, a comprehensive and coherent picture of changes in resting-state M/EEG activity in migraine is lacking so far.

Beyond providing insights into the pathophysiology of migraine, M/EEG might also help to develop clinically valuable biomarkers of migraine. According to the BEST (Biomarkers, EndpointS, and other Tools) framework [10], such biomarkers could serve different functions. *Diagnostic* biomarkers could support diagnosis and improve phenotyping and defining subtypes of migraine. *Monitoring* biomarkers would enable tracking disease trajectories and assessing treatment responses. *Predictive* biomarkers would be valuable in predicting treatment responses. *Prognostic* biomarkers might predict migraine attacks and disease courses. All these biomarkers would eventually help to optimize treatment and reduce suffering from migraine. Consequently, developing such biomarkers is a key area of current migraine research [11]. Resting-state electroencephalography (EEG) might be particularly suitable to establish migraine biomarkers since it is affordable, broadly available, potentially mobile, and scalable to large patient numbers. Magnetoencephalography (MEG) is closely related to EEG with even higher spatial and temporal resolution; however, it is more costly and less widely available.

Thus, resting-state M/EEG can further the understanding of the pathophysiology of migraine and serve to develop clinically valuable biomarkers. However, previous findings are inconsistent and have not yet been synthesized quantitatively. Hence, in this study, we aimed to summarize resting-state M/EEG findings in migraine through a systematic review and meta-analysis.

## 2 Methods

The study was conducted and is reported in accordance with the Preferred Reporting Items for Systematic Reviews and Meta-Analyses (PRISMA) guidelines [12]. The study protocol was registered on PROSPERO (Identifier *CRD42024550157*). Deduplication of records, title and abstract screening, full-text review, and data extraction were conducted using the Covidence software [13].

### 2.1 Search Strategy

Search strings comprised combinations of migraine and M/EEG, using Boolean operators and truncations. The complete search strings for all databases are detailed in the supplementary material. All databases were searched from their inception dates until May 27th, 2024. The databases MEDLINE, PubMed Central, and Bookshelf (through PubMed), Web of Science Core Collection (through Web of Science), and EMBASE (through Ovid) were searched. No language limit was applied. In addition, reference mining of recent reviews on M/EEG in migraine [5, 6, 14] and of included studies was performed. Furthermore, personal files were screened for relevant records.

### 2.2 Study Selection

Detailed inclusion and exclusion criteria are outlined in Table 1. In summary, we included peer-reviewed studies that used quantitative resting-state M/EEG to analyze brain activity in episodic or chronic migraine, either with or without aura. We excluded studies involving people with severe psychiatric or neurological comorbidities that could affect M/EEG activity, such as schizophrenia or multiple sclerosis. Additionally, we omitted rare subtypes of migraine (hemiplegic, retinal, brainstem aura, or pediatric migraine) with potentially different underlying pathophysiological mechanisms. Regarding associations of disease severity and M/EEG features, we included studies focusing on headache/attack frequency or headache intensity as measures of disease severity. These are regarded as the most established measures of disease severity in migraine studies [15] and were anticipated to be the most frequently reported.

**Table 1.**
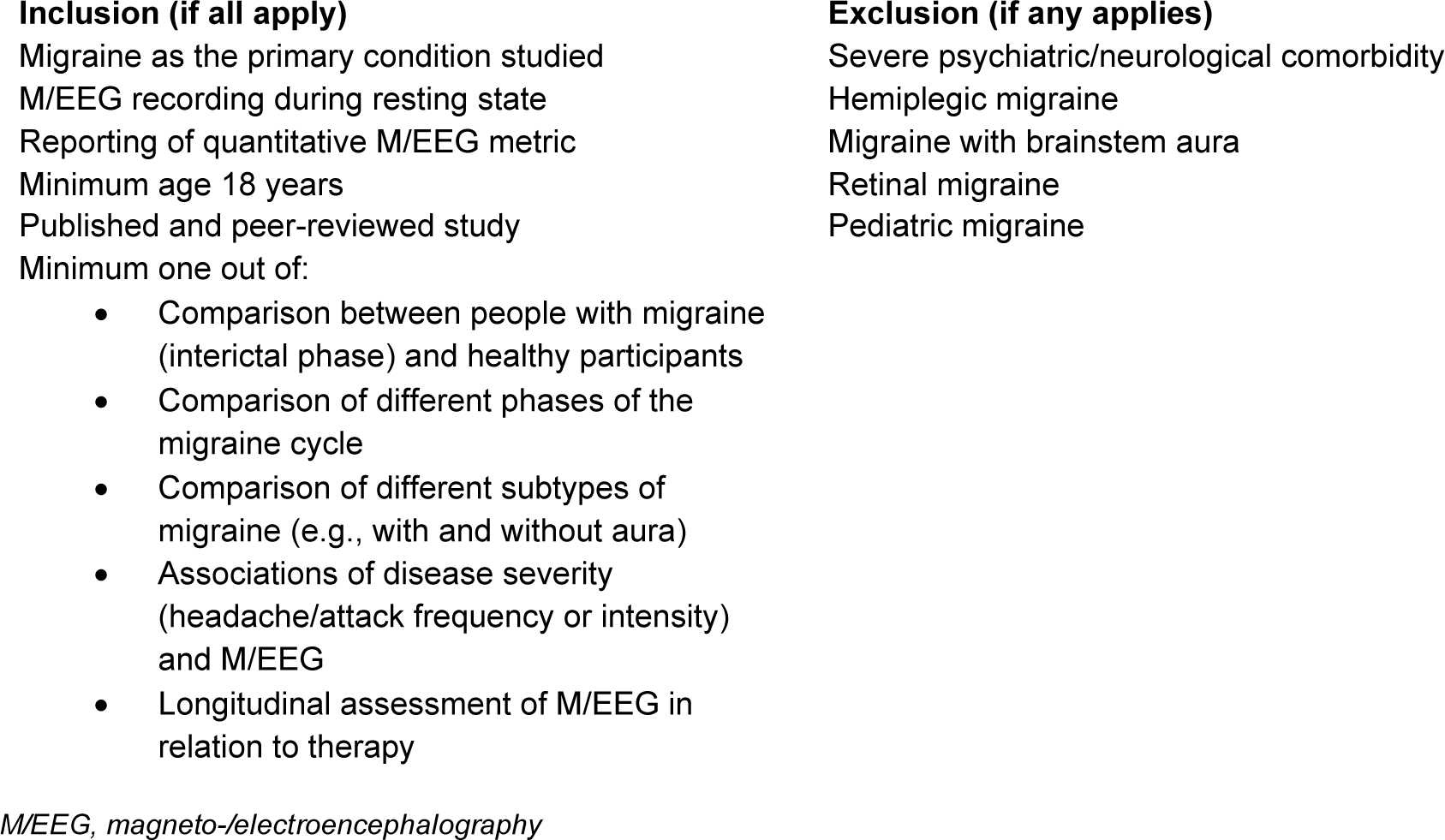
Inclusion and Exclusion Criteria.

### 2.3 Record screening, full-text review, and data extraction

Two authors independently screened titles and abstracts without knowing each other’s decisions. In case of disagreement, conflicts were discussed and resolved. The same procedure was followed for the full-text review. One author extracted data, and another author verified the results. Extracted data comprised general study information, details on participants (e.g., age, gender, migraine subtype), M/EEG recording specifications (e.g., electrode number, length of recording), and M/EEG outcome metrics (e.g., theta oscillatory power). For group comparisons of M/EEG features (t-tests and Mann-Whitney-U-tests), the following parameters were extracted for meta-analysis, if available: Means and standard deviations (SDs), t-values, U-values, and p-values. We followed recent recommendations [16] and implemented algebraic recalculation of means, SDs, and effect sizes if necessary. Data was extracted from figures whenever necessary and possible. We contacted study authors to retrieve statistics whenever algebraic recalculation was mathematically impossible.

If multiple comparisons were reported for one M/EEG feature (e.g., a study had analyzed several regions of interest for oscillatory theta power), the largest effect was selected for further analysis. If imprecise p-values for significant findings were reported (for example, “*p < 0.01*”), we used the closest decimal value for extraction (”*p = 0.009*”). If p-values for non-significant findings were reported (e.g., “*p > 0.05*”), we chose not to extract the nearest decimal because a valid approximation to the measured effect could not be ensured.

### 2.4 Data Synthesis

We used a multi-step approach for data synthesis, considering the quality and number of studies for each comparison (e.g., oscillatory theta power between people with migraine and healthy participants). For studies that analyzed peak alpha frequency (PAF), oscillatory power, or connectivity, we first used effect direction plots with vote-counting for semi-quantitative data synthesis. We interpreted comparisons with k>2 studies. Second, meta-analysis using R Version 4.1.2 [17] with the *metafor* package [18] was performed in case of k>4 suitable studies, following recent recommendations on the minimum number of studies needed for random-effect meta-analysis [19]. Random-effect models were chosen due to anticipated significant between-study heterogeneity in (i) M/EEG data acquisition and analysis and (ii) clinical characteristics of study participants. Heterogeneity was evaluated with Cochran’s *Q* (p < 0.05 indicating heterogeneity) and *I^2^*(values of 25%, 50%, and 75% representing low, moderate, and high heterogeneity, respectively). Funnel plots and Egger’s tests were used to assess publication bias. Due to the small sample sizes of studies, Hedges’ g was used to compare M/EEG features between groups. To that end, for studies using parametric statistical tests (t-tests), effect sizes were calculated directly from means and SDs, or p-values, and sample sizes. For studies using non-parametric tests (Mann-Whitney-U-tests), eta-squared was calculated as an effect size estimate [20] and converted to Hedges’ g using the *esc* package [21] in R. Narrative data synthesis was used for the remaining studies (i.e., other rarely used or less established M/EEG features and comparisons).

### 2.5 Risk of Bias and Quality Assessment

Risk of bias (RoB) and study quality were assessed using a modified version of the Newcastle– Ottawa Scale [22] for migraine and EEG studies (see supplemental material). This tool assesses the RoB and study quality in terms of “selection of study participants,” “comparability/confounders,” and “outcome data.” In the original version, stars are awarded for individual domains, whereas we rated items as “high” or “low” RoB for easier interpretation.

### 2.6 Missing Data and Full Texts

Corresponding authors were contacted up to two times via email to request missing data or inaccessible full texts. Data/full texts were considered unavailable if no reply was received two weeks after the second contact attempt.

## 3 Results

### 3.1 Study Selection

Database searches resulted in 2359 records after deduplication. Following title and abstract screening, 113 studies were identified. Finally, 27 studies [9, 23–48] were included. Fig. 1 shows the PRISMA flow diagram of study selection and exclusion reasons at different levels.

**Fig. 1.**
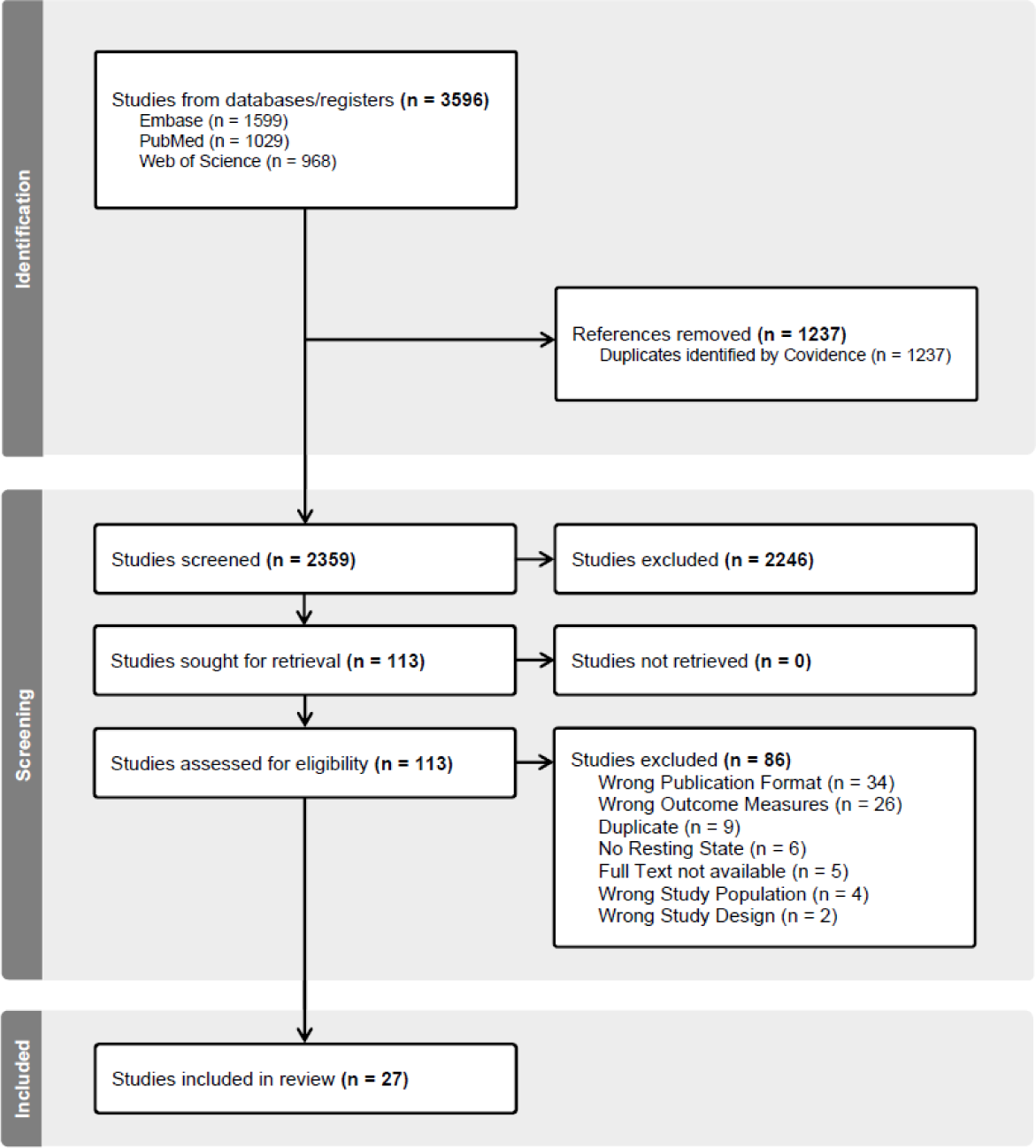
PRISMA flow diagram of study selection.

### 3.2 Study Characteristics

Fig. 2 and Fig. 3 summarize the main characteristics of the included studies. Regarding recording modality, most studies used EEG (*k = 22*). Regarding migraine type, most studies (*k = 11*) included both people with migraine with and without aura or did not differentiate between them (*k = 10*). Furthermore, most studies investigated people with episodic migraine (*k = 11*) or did not differentiate between episodic and chronic migraine (*k = 11*). Fig. 3 summarizes the performed analyses and investigated M/EEG features of studies. Most studies (*k = 20*) compared M/EEG recordings of people with migraine in the interictal phase to healthy participants. Five studies investigated differences between the interictal and the premonitory phase, and six studies analyzed correlations of M/EEG features with measures of disease severity. Three studies investigated differences between episodic and chronic migraine, and five studies between migraine with and without aura. Three studies compared M/EEG recordings of people with migraine during headache attacks or in the postictal phase with healthy participants. Three studies analyzed M/EEG features in longitudinal designs. The total sample sizes of included studies ranged from *n = 20* to *n = 215* with a mean total sample size of *n = 62.4* (*mean = 39.2 [range 10-150]* for people with migraine, *mean = 27.2 [range 10-66]* for healthy participants). Fig. S4 shows boxplots of study sample sizes for people with migraine, healthy participants, and total participants.

**Fig. 2.**
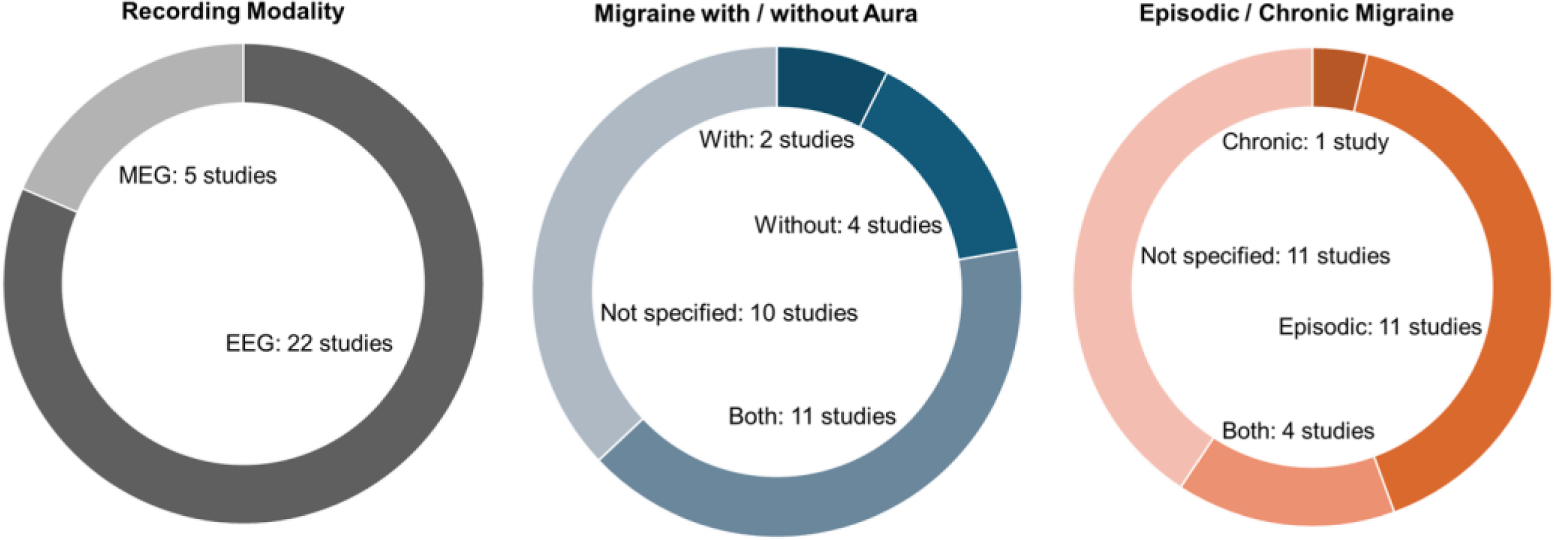
Overview of included studies.

**Fig. 3.**
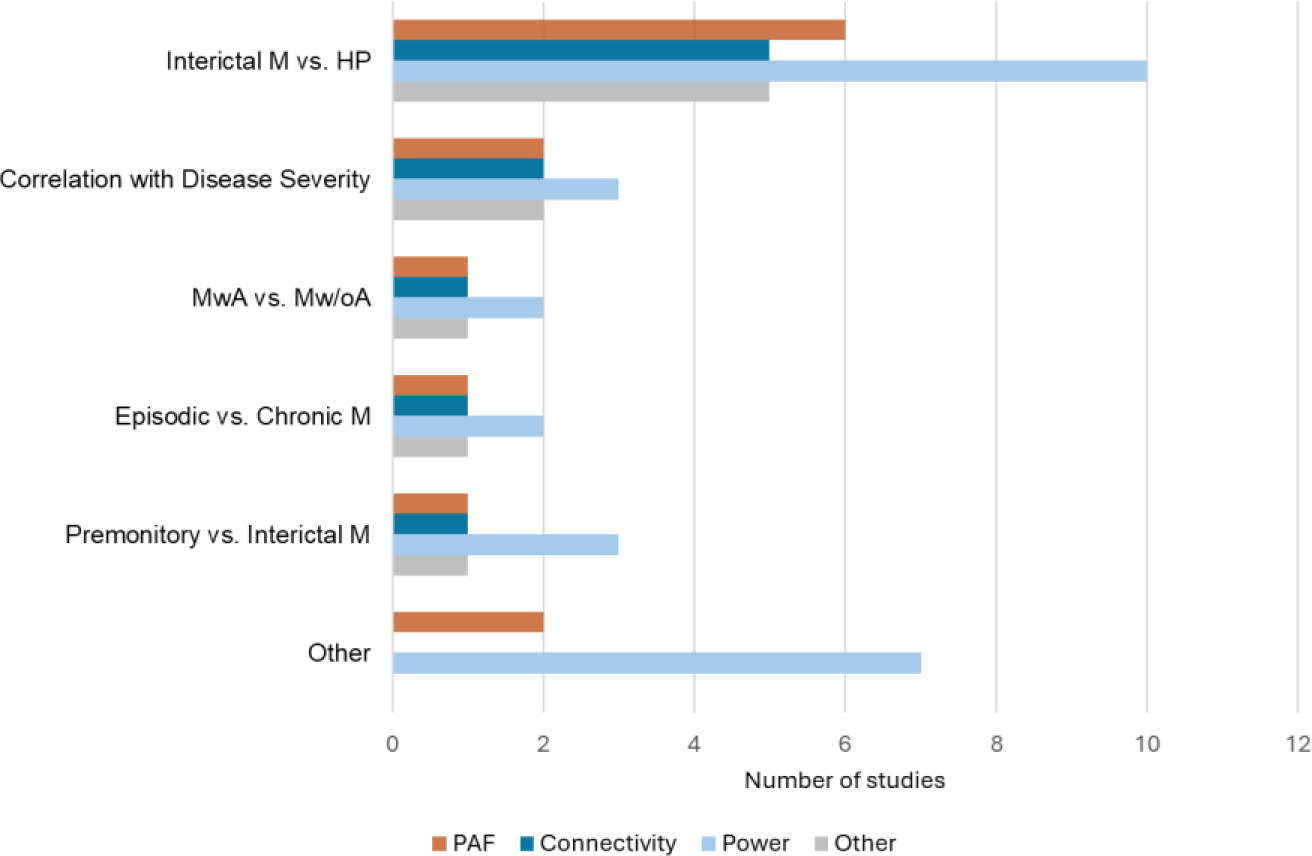
Number of included studies by comparison and analyzed M/EEG features. M=Migraine, HP=Healthy *Participants, MwA=Migraine with Aura, Mw/oA=Migraine without Aura, PAF=peak alpha frequency. Note that single studies occur multiple times for different comparisons*

### 3.3 Risk of Bias and Study Quality

A summary of the RoB assessment is presented in Fig. 4. Scoring of individual studies is presented in the supplementary material. In the domain “selection of study participants,” a significant RoB arose from “case representativeness,” as 41% of studies did not describe their sampling strategy. For the item “case definition,” RoB was low for most studies because diagnostic criteria were specified. For the items “matching of controls” and “definition of controls,” low RoB was obtained. In the domain “comparability/confounders,” RoB was high for most studies for the item “controlling for depression/anxiety”, because these frequent comorbidities of migraine were rarely controlled for, which could impact M/EEG activity. However, most studies controlled for medication use, which resulted in low RoB for the item “controlling for any other factor.” Regarding “outcome data,” RoB for the items “data acquisition/processing” and “appropriate statistical test” for most studies was low. However, most studies had a high RoB regarding the item “assessment of outcome” because preprocessing/analysis of M/EEG data was performed manually, and blinding to condition or clinical data was not explicitly stated.

**Fig. 4.**
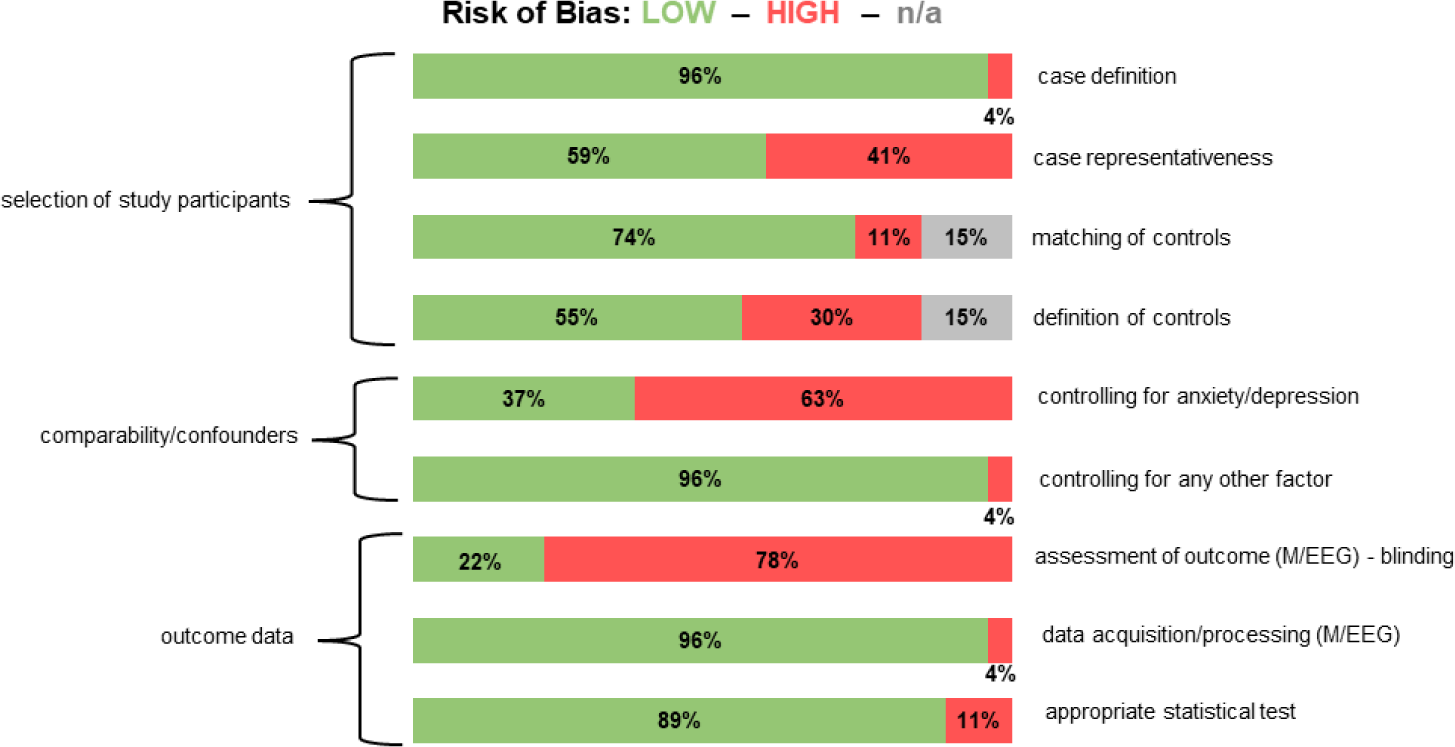
Overview of risk of bias assessment.

In summary, for most items (7/9), low RoB was found. The strongest RoB arose from a lack of controlling for comorbidities as potential confounders of M/EEG activity and from non-automatized or non-blinded M/EEG assessment.

### 3.4 Data Synthesis

#### 3.4.1 People with Migraine (Interictal Phase) vs. Healthy Participants

##### Vote Counting

Thirteen studies compared PAF or frequency-specific power between people with migraine in the interictal phase and healthy participants. Fig. 5 shows an effect direction plot of individual studies. For PAF, most studies (83%) found no differences between groups. Regarding delta power, most studies (75%) found no differences between groups. An equal number of studies found higher theta power and no differences in theta power (each 44%) between groups. Regarding alpha power, most studies (50%) found no differences between groups. For gamma power, an equal number of studies found higher values and no differences between groups (each 50%).

**Fig. 5.**
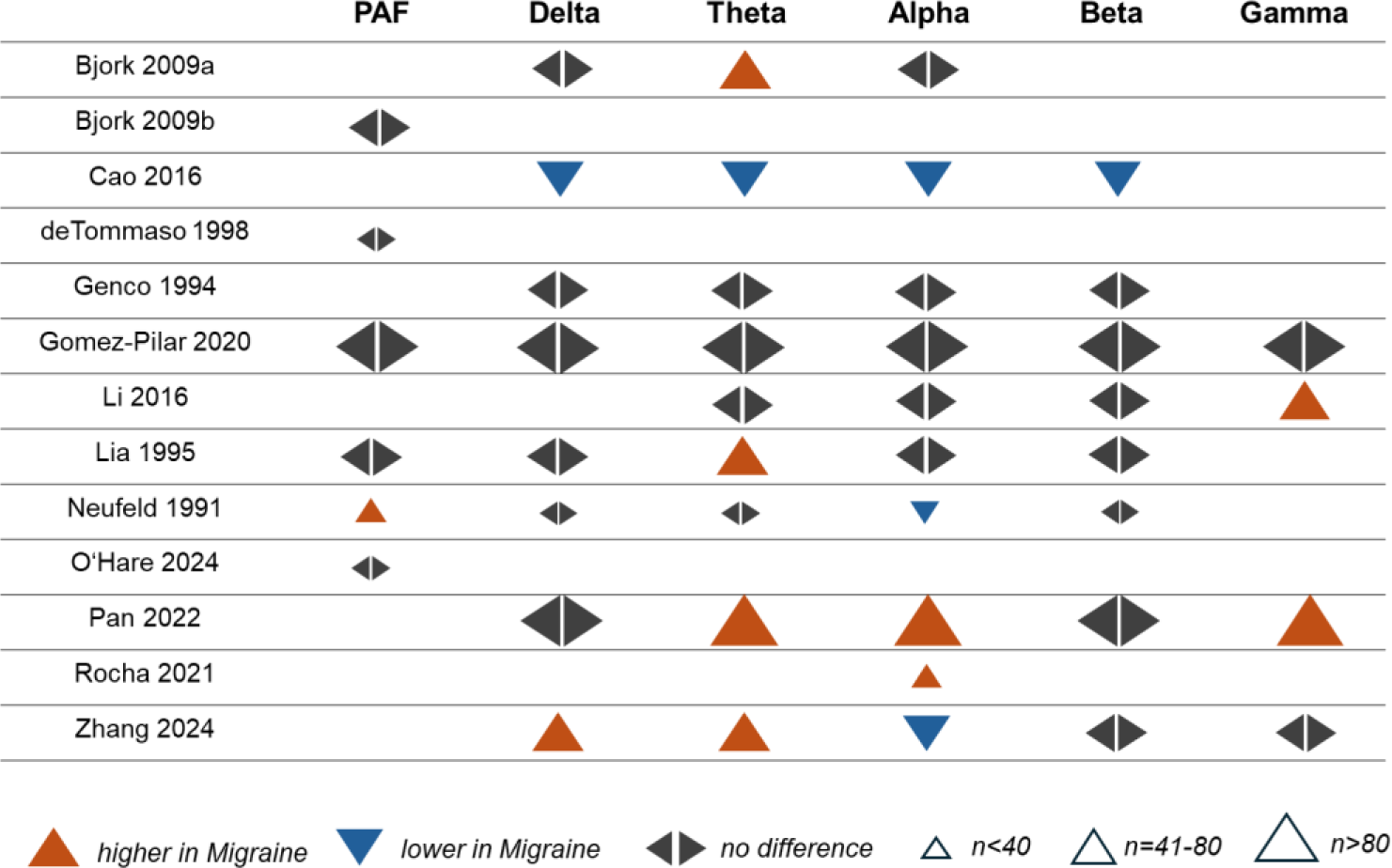
Effect direction plot for studies comparing peak alpha frequency and oscillatory power between people with migraine in the interictal phase and healthy participants. *PAF = peak alpha frequency, n = total sample size*.

In summary, vote counting revealed that most studies did not find differences in PAF or oscillatory power between people with migraine in the interictal phase and healthy participants. However, for theta and gamma power, the number of studies reporting higher values for people with migraine was the same as those reporting no differences. Moreover, the number of studies reporting higher values for theta power and gamma power was higher than those reporting lower values.

Five studies compared frequency-specific connectivity between people with migraine in the interictal phase and healthy participants. Fig. 6 shows an effect direction plot of individual studies. Regarding delta connectivity, most studies (75%) found no differences between groups. For theta, alpha, and beta connectivity, most studies (60%) found lower values in people with migraine. Regarding gamma connectivity, most studies (75%) found no differences between groups.

**Fig. 6.**
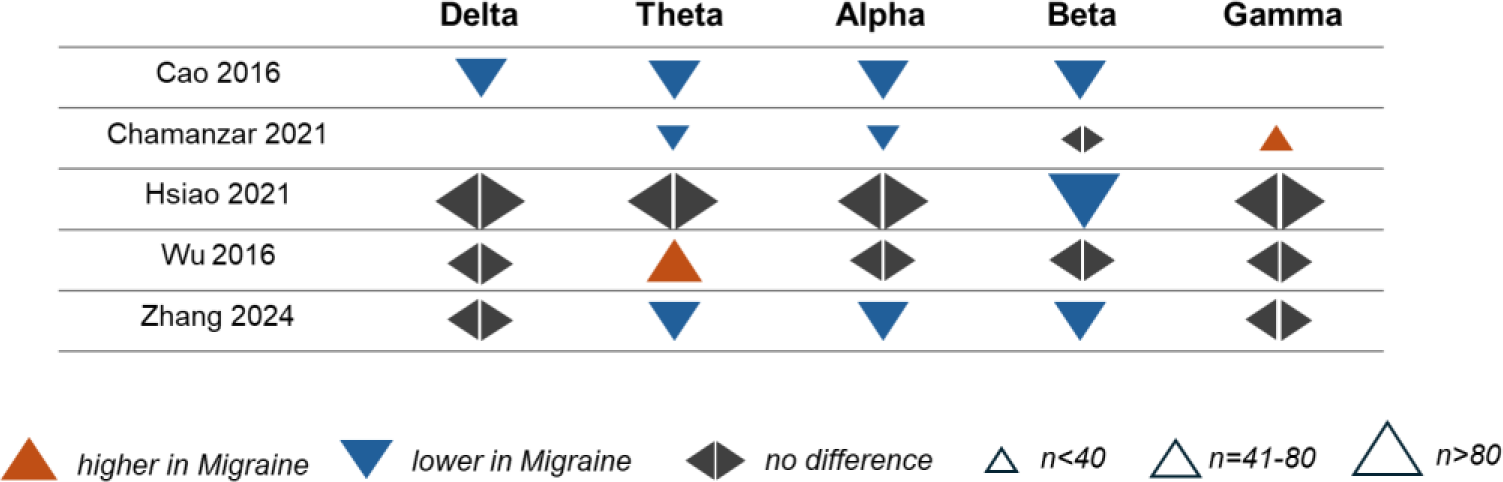
Effect direction plot for studies comparing connectivity between people with migraine in the interictal phase and healthy participants. *n = total sample size*

In summary, most studies found lower theta, alpha, and beta connectivity in people with migraine in the interictal phase compared to healthy participants. For delta and gamma connectivity, most studies found no differences between groups.

##### Meta-Analysis

Meta-analysis for group differences was performed for comparisons with k > 4 studies with necessary statistics available (*see 2.4, Data Synthesis*). Therefore, meta-analyses could be performed for delta, theta, and alpha power, and theta connectivity. Results are shown in Fig. 7. For delta power (*k = 5* studies comprising *n = 247* people with migraine and *n = 162* healthy participants), meta-analysis yielded non-significant differences between groups with a moderate degree of heterogeneity among studies (*Hedges’ g = 0.15, 95% CI -0.20-0.50, I^2^ = 64.9%, p [Q] = 0.038*). In contrast, for theta power (*k = 7* studies comprising *n = 299* people with migraine and *n = 214* healthy participants), we found a small to moderate effect for higher amplitudes in migraine with a moderate degree of heterogeneity among studies (*Hedges’ g = 0.38, 95% CI 0.02-0.74, I^2^ = 73.1%, p [Q] = 0.0034*). Regarding alpha power (*k = 8* studies comprising *n = 320* people with migraine and *n = 207* healthy participants), no group differences were found, while a high degree of heterogeneity among studies was observed (*Hedges’ g = -0.04, 95% CI -0.53-0.46, I^2^ = 85.9%, p [Q] < 0.0001*). Although funnel plots suggested a certain asymmetry, Egger’s tests did not yield significant results, arguing against publication bias (see Fig. S1).

**Fig. 7.**
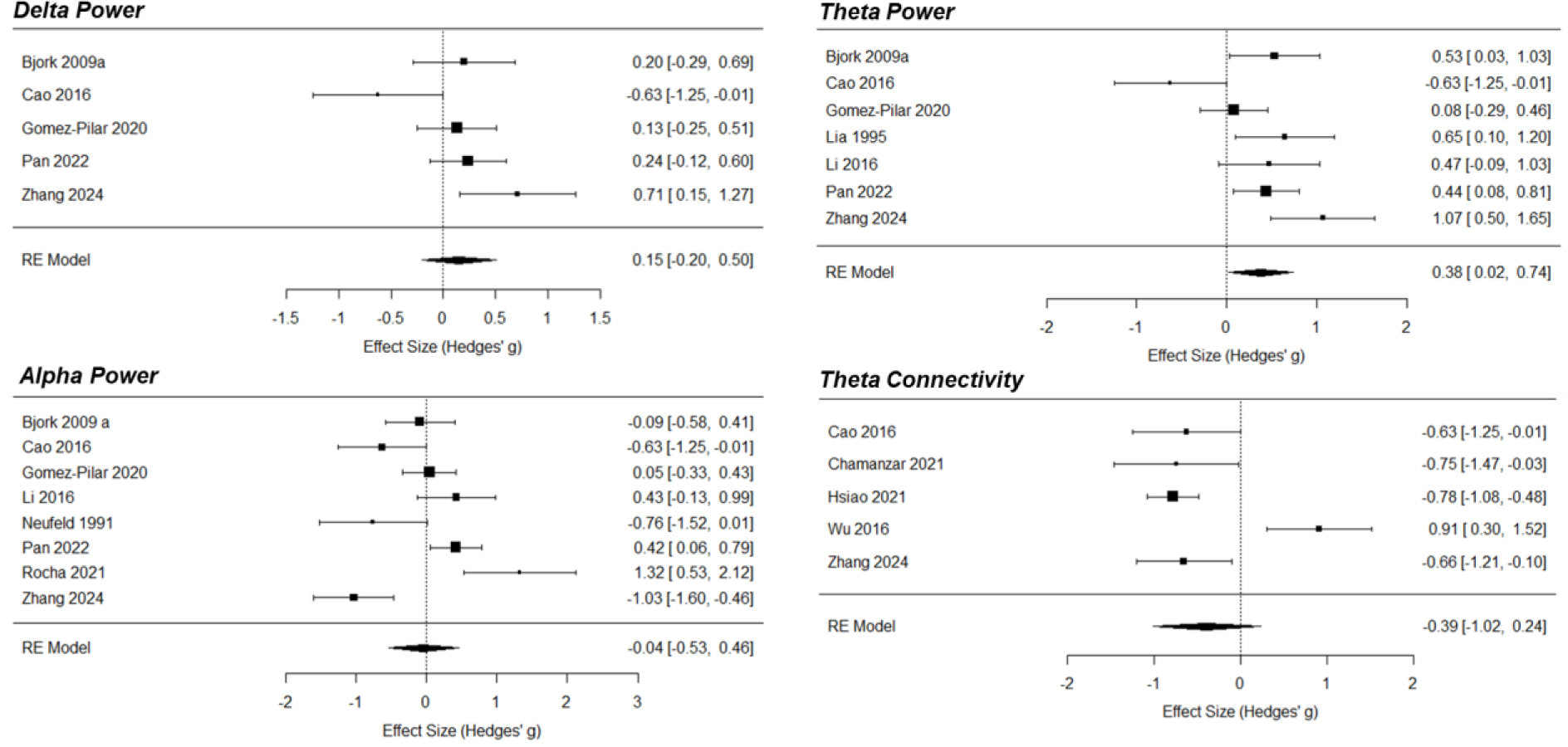
Forest plots of meta-analyses for comparison of delta, theta, and alpha power and theta connectivity between people with migraine in the interictal phase and healthy participants. *Note that Hsiao et al. (2021) did not report group differences in the primary study after correcting for multiple comparisons. However, a significant effect was observed in the meta-analysis based on the study’s primary data (means and standard deviations)*.

Meta-analysis for group differences in theta connectivity (*k = 5* suitable studies comprising *n = 242* people with migraine and *n = 148* healthy participants) yielded non-significant differences between groups with a high degree of heterogeneity among studies (*Hedges’ g = -0.39, 95% CI -1.02-0.24, I^2^ = 86.65%, p [Q] < 0.001*). The funnel plot and Egger’s test (*p > 0.5*) did not show evidence for publication bias (see Fig. S1).

In summary, meta-analysis showed higher theta power in people with migraine in the interictal phase compared to healthy participants. No significant differences were observed for delta and alpha power, as well as for theta connectivity.

#### 3.4.2 Interictal Phase vs. Premonitory Phase

Four studies compared PAF or frequency-specific power between the premonitory and interictal phases. Fig. 8 shows an effect direction plot of individual studies. Regarding delta power, most studies (67%) found higher values in the premonitory phase. For both theta and alpha power, most studies (each 67%) found no differences. Regarding beta power, most studies (67%) found higher values in the premonitory phase. For PAF, gamma power, and connectivity, the number of included studies was too low for meaningful vote counting.

**Fig. 8.**
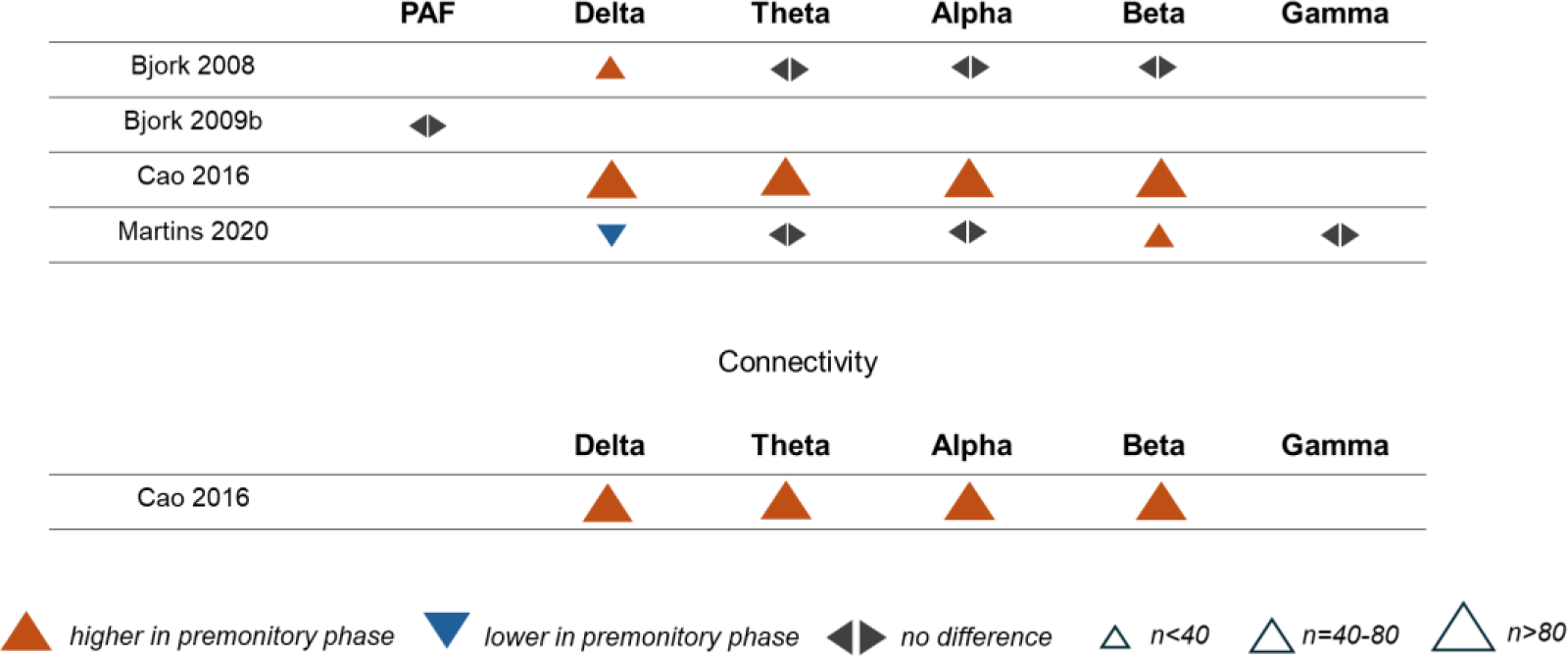
Effect direction plot for studies comparing peak alpha frequency, oscillatory power, or connectivity between the premonitory phase and the interictal phase. *PAF = peak alpha frequency, n = total sample size*.

In summary, most studies found increases in delta and beta power in the premonitory phase but no changes in theta and alpha power.

#### 3.4.3 Correlations with Disease Severity

Four studies investigated correlations between the frequency of attack/headache days and PAF or frequency-specific power. Two studies investigated correlations between the frequency of attack/headache days and frequency-specific connectivity, and two studies investigated correlations between headache intensity and frequency-specific power. Effect direction plots for included studies are shown in Fig. 9. With three exceptions, non-significant correlations were found.

**Fig. 9.**
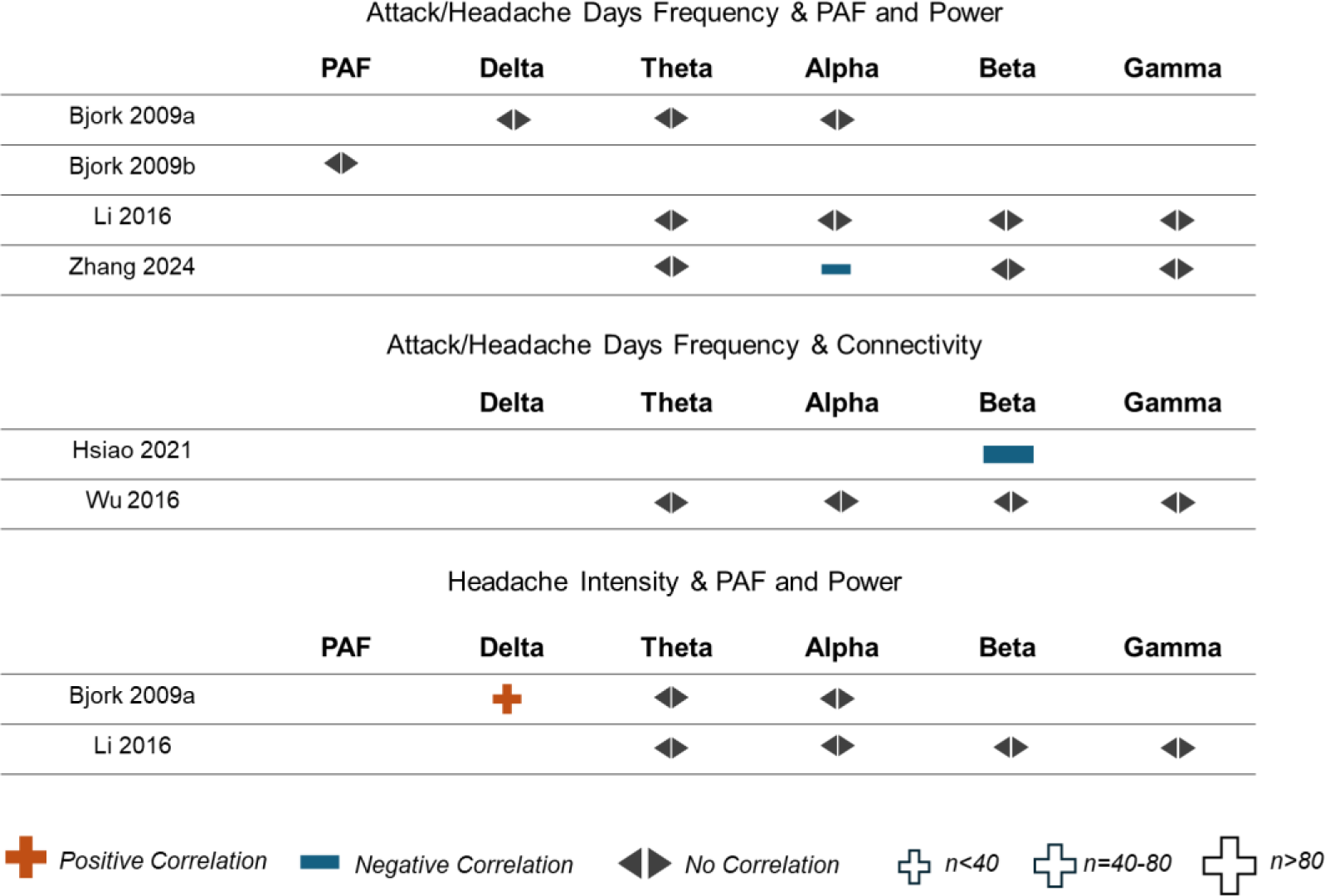
Effect direction plot for studies correlating measures of disease severity with peak alpha frequency, oscillatory power, or connectivity. *PAF = peak alpha frequency, n = total sample size*.

#### 3.4.4 Episodic vs. Chronic Migraine

Two studies compared PAF or frequency-specific power, and two studies compared frequency-specific connectivity between people with episodic and chronic migraine. Hence, the number of included studies was too low for meaningful vote counting. An effect direction plot of included studies is shown in Fig. S2.

#### 3.4.5 Migraine with vs. without Aura

Three studies compared PAF, two studies compared frequency-specific power, and one study compared frequency-specific connectivity between people with migraine with and without aura. An effect direction plot of included studies is shown in Fig. S3. Regarding PAF, all studies found no differences between groups. For power and connectivity, the number of included studies was too low for meaningful vote counting.

#### 3.4.6 Other Studies (Narrative Data Synthesis)

Two studies compared oscillatory power between people with migraine during a headache attack and healthy participants. One study found no differences regarding delta, theta, alpha, and beta power [32]. The other found higher theta and lower alpha power in people with migraine [29]. One study compared oscillatory power between people with migraine in the postictal phase and healthy participants and found lower delta, theta, and beta power in people with migraine [27].

Three studies [23, 34, 36] used microstate analysis to evaluate differences between people with migraine in the interictal phase and healthy participants. Microstate analysis characterizes EEG activity as a sequence of stable states lasting tens of milliseconds before rapidly transitioning to another. Typically, resting-state EEG activity is represented by 4 to 6 microstates, highly consistent across individuals, and labeled with letters. All three studies identified a more frequent occurrence of microstate B in people with migraine, but two found these changes only in the subsample without aura [34, 36]. Differences regarding other microstates were inconsistent between studies.

One study [45] compared EEG complexity (“fuzzy entropy”) between people with migraine and healthy participants and between different phases of the migraine cycle. The authors found an increase of EEG complexity in the premonitory phase and a decrease in the interictal phase compared to healthy participants.

Three studies investigated oscillatory power in the context of longitudinal study designs. One study found higher baseline alpha power in responders compared to non-responders to prophylactic treatment with flunarizine [28]. However, no corresponding changes in alpha power paralleling treatment effects were found. Another study found decreases in delta, alpha, and beta power along with treatment response to various prophylactic agents [37]. Yet another study found a decrease in alpha power after 12 sessions of transcranial direct current stimulation [26], but no clinical measures of treatment effects were provided.

## 4 Discussion

### 4.1. Main findings

The present systematic review and meta-analysis synthesizes evidence on M/EEG findings in migraine. The meta-analysis showed increased power of theta activity in people with migraine in the interictal phase compared to healthy participants. Based on vote counting, we found evidence for lower connectivity at alpha and beta frequencies in people with migraine in the interictal phase. In other frequency bands or PAF, we found no differences between groups.

Increased theta power in people with migraine is consistent with findings in chronic pain [49], pathological fatigue [50], and various psychiatric disorders, including mood disorders [51]. Thus, increased theta activity appears not to be specific for migraine but to represent a shared feature of different pain conditions and neuropsychiatric disorders. The specific clinical correlate of increased theta activity remains unknown. The variety of conditions associated with it suggests that it represents a general feature of different brain disorders, such as a heightened vulnerability or a shared symptom, such as depressed mood or suffering. Large-scale studies considering neuropsychiatric symptoms with disease-spanning clinical assessments might help to understand the functional significance of increased theta power in migraine and other brain disorders.

In addition, we found evidence for lower alpha and beta connectivity in people with migraine in the interictal phase compared to healthy participants. This finding was based on vote counting, as meta-analysis was feasible for theta connectivity only. Here, a small to moderate effect for lower theta connectivity in people with migraine failed to reach statistical significance. Brain connectivity plays an essential role in pain processing [52–54], and alterations have been found in different neurological and psychiatric conditions [55]. For migraine as a brain network disorder [4], changes in connectivity patterns are highly plausible and should be investigated further.

While the features above may serve as diagnostic biomarkers, the limited number of studies did not allow for drawing valid conclusions regarding predictive/prognostic biomarkers. Only a few (*k = 4* for oscillatory power, *k = 1* for connectivity) studies investigated M/EEG variations across the migraine cycle and compared people with migraine in the premonitory and the interictal phase. Some evidence for increased delta and beta power was found. However, objectively identifying biomarkers for the premonitory phase, especially for patients not regularly experiencing prodromal symptoms, would be clinically highly valuable. Predicting treatment response is considered a key challenge in current migraine research[11], and a non-invasive, easy-to-use technology like EEG offers untapped potential. However, only one study [28] used baseline M/EEG as a predictor for treatment response (and found higher baseline alpha power in later respondents to prophylaxis with flunarizine).

### 4.2. Limitations

The included studies yielded limitations of the present systematic review and meta-analysis. First, meta-analysis was feasible only for delta, theta, and alpha power and for theta connectivity regarding comparisons between people with migraine in the interictal phase and healthy participants. Other comparisons (e.g., connectivity in other frequency bands or M/EEG changes in the premonitory phase) or analysis of associations with disease severity had to be based solely on vote counting. Similarly, detailed analyses regarding episodic and chronic migraine or migraine with and without aura were not feasible. Second, considerable methodological heterogeneity between studies existed. For instance, some studies investigated global oscillatory power [39, 40], while others focused on regions of interest, e.g., occipital electrodes [26, 31]. Due to the low number of studies, these studies were pooled. This might obscure features and biomarkers specific to certain brain regions or networks. Third, the sample sizes of studies were mostly low. The mean total sample size of included studies was *n = 62.4,* while most studies investigated between-group comparisons. Sample size calculation indicates that for detecting a medium effect with a power of 80% (two-tailed t-test, *alpha = 0.05, 1-beta = 0.8*), a total sample size of *n = 128* would be necessary [56]. Thus, most studies were underpowered, which might imply a low sensitivity and a high risk of false positive findings [57].

### 4.3 Outlook and Recommendations

The present findings can help to guide future M/EEG studies in migraine. The following aspects might warrant further investigation.

First, most studies focused on band-specific power of brain activity. However, other M/EEG features might also be investigated more thoroughly. Connectivity measures are particularly promising as functional imaging studies have revealed dysfunction of connectivity within and across brain networks in migraine [4]. However, only a few M/EEG studies have assessed connectivity in migraine so far. Imbalances between cerebral excitation and inhibition might be another promising target for M/EEG studies. Such an imbalance has been indicated by a hypersensitivity to sensory stimuli [3, 58] and evoked potential, transcranial magnetic stimulation, and functional MRI studies [4, 6]. New approaches now allow the quantification of the relationship between excitation and inhibition based on M/EEG recordings [59]. For instance, the aperiodic component of the EEG signal [60] is a promising feature, which is altered in various neuropsychiatric disorders [61] but has not been investigated in migraine so far.

Second, most studies focused on features and comparisons which allow for developing diagnostic biomarkers. However, other biomarker types can also be relevant. For instance, predicting responses to a specific treatment would be highly relevant. Clinical information alone often does not allow for the prediction of treatment responses, and even for treatments with monoclonal antibodies targeting CGRP, a considerable number of non-responders exist [62]. Moreover, predicting migraine attacks would be clinically valuable. Such a prediction could allow for preventing progression to the headache phase, e.g., by gepants [63], long-acting triptans [64], or neuromodulatory procedures. Mobile, user-friendly EEG systems are feasible and practical solutions for such situations (13).

Third, most studies did not control for relevant comorbidities like depression. Thus, the specificity of many of the findings remains unclear. Future studies on migraine should assess comorbidities in more detail. This is important both for the pathophysiological understanding and for the evaluation of brain-based biomarkers of migraine. Larger, multivariate studies should assess comorbidities and other pain conditions and relate them to M/EEG alterations.

Fourth, recording and analysis procedures were not standardized. However, the robustness and replicability of findings can only be achieved by high standards in conducting and reporting data acquisition and analysis [65, 66]. Moreover, collaborative approaches and data-sharing initiatives hold untapped potential in M/EEG-based biomarker research in migraine. The recently established ENIGMA Chronic Pain Working Group is an example of such an initiative [67].

### 4.4 Conclusions

The present systematic review and meta-analysis of resting-state M/EEG findings in people with migraine revealed higher power of brain activity at theta frequencies compared to healthy participants. Furthermore, we found evidence for lower alpha and beta connectivity in people with migraine in the interictal phase. The most substantial risk of bias arose from a lack of controlling for comorbidities and non-automatized or non-blinded M/EEG assessments. Together, these findings can guide future M/EEG studies on migraine pathophysiology and brain-based biomarkers, which should consider comorbidities, address new EEG measures, and aim for standardized, collaborative approaches.

## Supporting information

Supplemental Material_all

## Declarations

### Ethics approval and consent to participate

Not applicable (no primary data was generated for this study)

### Consent for publication

Not applicable (no primary data was generated for this study)

### Availability of data and materials

All search strings used for literature search and R-code for meta-analyses can be found in the supplementary material. No primary data was generated for this study.

### Authors’ contributions

**PTZ** Conceptualization, Literature Search, Data Analysis, Writing, Review & Editing; **HH** Literature Search, Data Analysis, Review & Editing; **EM** Data Analysis, Review & Editing; **MP** Conceptualization, Literature Search, Data Analysis, Writing, Review & Editing.

## Acknowledgments

The authors used Grammarly to improve language and readability while preparing this manuscript. The authors reviewed and edited the content as needed and took full responsibility for the content of the work.

