## Supplemental Material_all for "Resting-State Electroencephalography and Magnetoencephalography in Migraine – A Systematic Review and Meta-Analysis"

Supplementary Material

**Fig. S1** Funnel plots and Egger's Test results for publication bias of meta-analyses

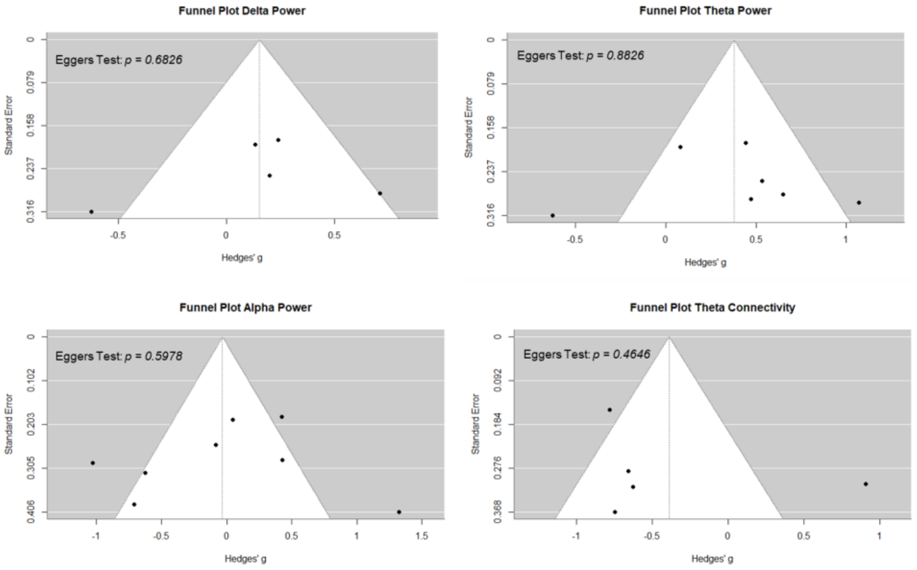


**Fig. S2** Effect direction plot for comparing PAF, oscillatory power, or connectivity between episodic and chronic migraine

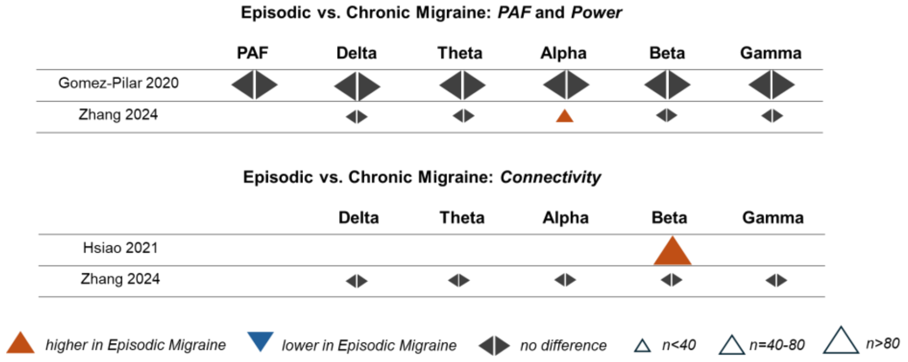


**Fig. S3** Effect direction plot for comparing PAF, oscillatory power, or connectivity between migraine with and without aura

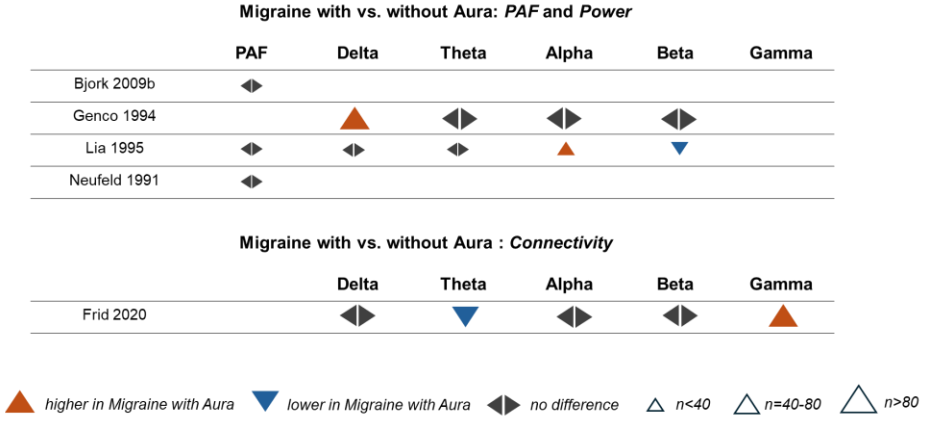

**Fig. S4** Sample size of individual studies

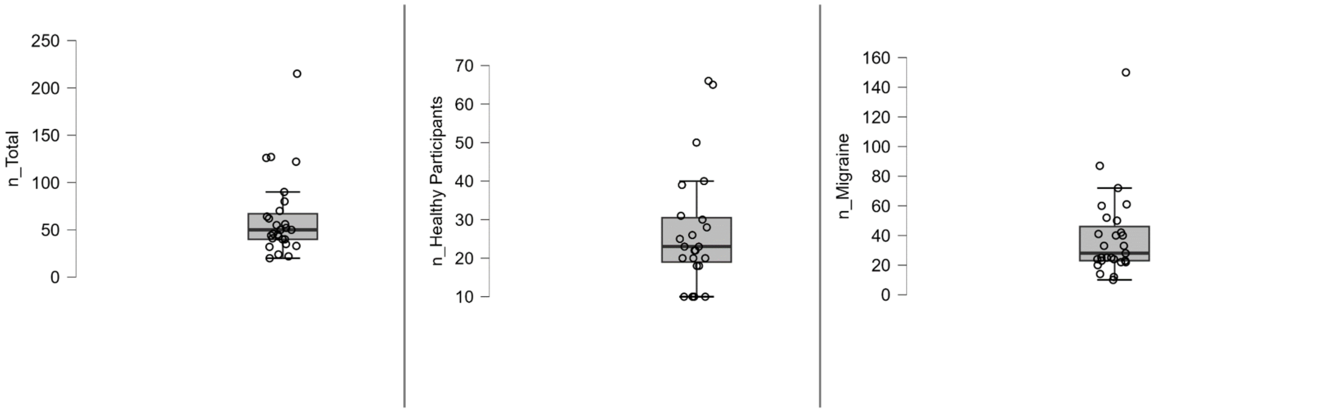


**Table S1** Individual scoring of Risk of Bias Assessment for included studies

| Study ID | Title | Selection: Case Definition | Selection: Representativeness | Selection: Controls | Selection: Definition of Controls | Comparability: Anxiety/Depression | Comparability: Any other factor | Outcome: Independent EEG preprocessing /analysis | Outcome: Description of EEG measurement | Outcome: Statistical test |
|---|---|---|---|---|---|---|---|---|---|---|
| Bjørk 2008 | Quantitative EEG power and asymmetry increase 36 h before a migraine attack. | Low RoB | Low RoB | | | High RoB | Low RoB | High RoB | Low RoB | Low RoB |
| Bjørk 2009 b | The occipital alpha rhythm related to the "migraine cycle" and headache burden: a blinded, controlled longitudinal study. | Low RoB | Low RoB | Low RoB | Low RoB | High RoB | Low RoB | High RoB | Low RoB | Low RoB |
| Bjørk 2009 a | Interictal quantitative EEG in migraine: a blinded controlled study. | Low RoB | Low RoB | Low RoB | Low RoB | High RoB | Low RoB | High RoB | Low RoB | Low RoB |
| Cao 2016 | Resting-state EEG power and coherence vary between migraine phases. | Low RoB | Low RoB | Low RoB | Low RoB | Low RoB | Low RoB | High RoB | Low RoB | High RoB |
| Cao 2018 | Exploring resting-state EEG complexity before migraine attacks. | Low RoB | High RoB | Low RoB | Low RoB | Low RoB | Low RoB | Low RoB | Low RoB | Low RoB |
| Chamanzar 2021 | Abnormalities in cortical pattern of coherence in migraine detected using ultra high-density EEG. | Low RoB | High RoB | Low RoB | Low RoB | High RoB | Low RoB | High RoB | Low RoB | Low RoB |
| deTommaso 1998 | EEG spectral analysis in migraine without aura attacks. | Low RoB | Low RoB | Low RoB | Low RoB | High RoB | Low RoB | High RoB | Low RoB | Low RoB |
| Frid 2020 | A Biomarker for Discriminating Between Migraine With and Without Aura: Machine Learning on Functional Connectivity on Resting-State EEGs. | Low RoB | High RoB | | | High RoB | Low RoB | Low RoB | Low RoB | Low RoB |
| Genco 1994 | EEG features in juvenile migraine: topographic analysis of spontaneous and visual evoked brain electrical activity: a comparison with adult migraine. | Low RoB | High RoB | Low RoB | High RoB | High RoB | Low RoB | High RoB | High RoB | High RoB |
| Gomez-Pilar 2020 | Exploring EEG Spectral Patterns in Episodic and Chronic Migraine During the Interictal State: Determining Frequencies of Interest in the Resting State. | Low RoB | Low RoB | Low RoB | High RoB | Low RoB | Low RoB | High RoB | Low RoB | Low RoB |
| Hsiao 2021 | Migraine chronification is associated with beta-band connectivity within the pain-related cortical regions: a magnetoencephalographic study. | Low RoB | High RoB | Low RoB | Low RoB | Low RoB | Low RoB | Low RoB | Low RoB | Low RoB |
| Kim 2023 | Quantitative electroencephalography as a potential biomarker in migraine. | Low RoB | Low RoB | | | High RoB | High RoB | High RoB | Low RoB | Low RoB |
| Lei 2023 | Resting-state electroencephalography microstate dynamics altered in patients with migraine with and without aura-A pilot study. | Low RoB | Low RoB | Low RoB | Low RoB | High RoB | Low RoB | High RoB | Low RoB | Low RoB |
| Li 2016 | Abnormal resting-state brain activity in headache-free migraine patients: A magnetoencephalography study. | Low RoB | High RoB | Low RoB | High RoB | High RoB | Low RoB | Low RoB | Low RoB | Low RoB |
| Li 2022 | Abnormalities in resting-state EEG microstates are a vulnerability marker of migraine. | Low RoB | High RoB | Low RoB | Low RoB | Low RoB | Low RoB | High RoB | Low RoB | Low RoB |
| LIA 1995 | COMPUTERIZED EEG-ANALYSIS IN MIGRAINE PATIENTS | Low RoB | High RoB | High RoB | Low RoB | High RoB | Low RoB | High RoB | Low RoB | Low RoB |
| Liu 2015 | Resting state brain activity in patients with migraine: a magnetoencephalography study. | Low RoB | High RoB | Low RoB | Low RoB | High RoB | Low RoB | High RoB | Low RoB | Low RoB |
| Martins 2020 | Brain state monitoring for the future prediction of migraine attacks. | Low RoB | High RoB | | | High RoB | Low RoB | High RoB | Low RoB | Low RoB |
| Neufeld 1991 | EEG and topographic frequency analysis in common and classic migraine. | Low RoB | Low RoB | High RoB | High RoB | High RoB | Low RoB | High RoB | Low RoB | High RoB |
| O'Hare 2024 | No Evidence of Cross-Orientation Suppression Differences in Migraine with Aura Compared to Healthy Controls. | High RoB | Low RoB | High RoB | High RoB | High RoB | Low RoB | Low RoB | Low RoB | Low RoB |
| Ojha 2024 | Resting-state Quantitative EEG Spectral Patterns in Migraine During Ictal Phase Reveal Deviant Brain Oscillations: | Low RoB | Low RoB | Low RoB | Low RoB | Low RoB | Low RoB | High RoB | Low RoB | Low RoB |

| | | | | | | | | | | |
|---|---|---|---|---|---|---|---|---|---|---|
| | <i>Potential Role of Density Spectral Array</i> | | | | | | | | | |
| Pan 2022 | Resting-state occipital alpha power is associated with treatment outcome in patients with chronic migraine. | Low RoB | Low RoB | Low RoB | Low RoB | Low RoB | Low RoB | High RoB | Low RoB | Low RoB |
| Pelayo-González 2023 | Quantitative Electroencephalographic Analysis in Women with Migraine during the Luteal Phase | Low RoB | Low RoB | Low RoB | Low RoB | Low RoB | High RoB | Low RoB | Low RoB | Low RoB |
| Rocha 2021 | Could cathodal transcranial direct current stimulation modulate the power spectral density of alpha-band in migrainous occipital lobe? | Low RoB | Low RoB | Low RoB | High RoB | High RoB | Low RoB | High RoB | Low RoB | Low RoB |
| Wu 2016 | Multi-frequency analysis of brain connectivity networks in migraineurs: a magnetoencephalography study. | Low RoB | High RoB | Low RoB | High RoB | High RoB | Low RoB | High RoB | Low RoB | Low RoB |
| Zhang 2024 | Altered Neuromagnetic Activity in the Default Mode Network in Migraine and Its Subgroups (Episodic Migraine and Chronic Migraine). | Low RoB | Low RoB | Low RoB | High RoB | Low RoB | Low RoB | High RoB | Low RoB | Low RoB |
| Zhou 2023 | Spatio-temporal dynamics of resting-state brain networks are associated with migraine disability. | Low RoB | Low RoB | Low RoB | Low RoB | Low RoB | Low RoB | Low RoB | Low RoB | Low RoB |

**S1** Newcastle-Ottawa quality assessment scale adapted for cross-sectional/ case-control studies. Adaption for M/EEG studies in Migraine

Note: A study can be awarded a maximum of one star for each numbered item within the Selection and Exposure categories. A maximum of two stars can be given for Comparability. Items are rated as *high* (indicating negative study quality) or *low* (indicating positive study quality) Risk of Bias.

**Selection (Maximum 4 stars)**

1) Is the case definition (migraine) adequate?
a) yes, with independent validation (eg. self reported doctor's diagnosis, reference to primary medical record source,
international criteria) **Low ROB**
b) no, based on self reports
c) no description

2) Representativeness of the cases
a) Truly representative of the average in the target population. (all subjects or random sampling) **Low ROB**
b) Somewhat representative of the average in the target population. (non-random sampling) **Low ROB**
c) Selected group of users
d) No description of the sampling strategy

3) Matching/Selection of Controls
a) Matched by sex and/or age **Low ROB**
b) Matched by other factor
c) No description

4) Definition of Controls
a) No history of migraine and no current severe neurological or psychiatric diagnosis **Low ROB**
b) No description of source

**Comparability (Maximum 2 stars)**
1) Comparability of cases and controls on the basis of the design or analysis
a) The study controls for the most important factor (anxiety/depression). **Low ROB**
b) The study controls for any additional factor (eg. age, medication usage, duration of migraine) **Low ROB**

**Outcome (Maximum 3 stars)**
1) Assessment of outcome (EEG findings)
a) Independent blind assessment **Low ROB**

b) Clear description of method for EEG data acquisition (number and placement of electrodes, equipment, sample
rating) and data processing (how the parameters were extracted from the EEG data) **Low ROB**
c) No description of method for EEG data acquisition and/or data processing

2) Statistical test
a) The statistical test used to analyze the data is clearly described and appropriate, and the measurement of the
association is presented, including confidence intervals/effect size and the probability level (p value).
**Low ROB**
b) The statistical test is not appropriate, not described or incomplete.

**S2** Search Strings

**MEDLINE, PubMed Central and Bookshelf (via PubMed)**
migrain* [Title/Abstract] AND (qEEG [Title/Abstract] OR EEG [Title/Abstract] OR electroencephalogr* [Title/Abstract] OR MEG [Title/Abstract] OR magnetoencephalogr* )

**Web of Science Core Collection (via Web of Science)**
migrain* AND (qEEG OR EEG OR electroencephalogr* OR MEG OR magnetoencephalogr*)
+Topic (Abstract/Title)

**EMBASE (via OVID)**
(migrain* AND (qEEG OR EEG OR electroencephalogr* OR MEG OR magnetoencephalogr*)).ab,ti.

**S3** Code (R Version 4.1.2)

**Calculation of hedges g for MWU-Tests**
```
library(esc)
# Given p-value
p_value <- 0.xyz
# Given total N (n1+2)
n1 <- x
n2 <- y
N <- x+y
# Calculate the z-score
z_score <- qnorm(p_value / 2, lower.tail = FALSE)
# Calculate eta-squared
eta_squared <- (z_score^2) / (N)
# convert to cohens d based on doi:10.1037/a0024338
d <- (2*sqrt(eta_squared))/(sqrt(1-eta_squared))
# convert cohens d to hedges g
totaln <- N
hedges_g(d, totaln)
```

**RE-Meta-Analysis**
```
library(metafor)
library(readxl)
# Specify the path to your Excel file
file_path <- read_excel("Path")
```

```
# Load the data from the Excel file
data <- read_excel("Path")
# Calculate the variances for Hedges' g
data$var_g <- (data$n1 + data$n2) / (data$n1 * data$n2) + (data$hedges_g^2 / (2 * (data$n1
+ data$n2)))
# Assign study labels to studies and data
study = c("Study1”)
study_labels <-data$study
# Conduct the meta-analysis
res <- rma(yi = hedges_g, vi = var_g, data = data, method = "REML")
# Print the results
print(res)
```

**Create a forest plot**

```
forest(res, slab = study_labels,  xlab = "Effect Size (Hedges' g)")
```

**Create a funnel plot**

```
funnel(res, main="Funnel Plot Theta Power", xlab="Hedges' g", ylab="Standard Error")
```

**Conduct Egger’s test**

```
egger_test <- regtest(res, model = "lm")
print(egger_test)
```
